# Characterizing the Adult and Larval Transcriptome of the Multicolored Asian Lady Beetle, *Harmonia axyridis*

**DOI:** 10.1101/034462

**Authors:** Lindsay Havens, Matthew MacManes

## Abstract

The reasons for the evolution and maintenance of striking visual phenotypes are as widespread as the species that display these phenotypes. While study systems such as *Heliconius* and *Dendrobatidae* have been well characterized and provide critical information about the evolution of these traits, a breadth of new study systems, in which the phenotype of interest can be easily manipulated and quantified, are essential for gaining a more general understanding of these specific evolutionary processes. One such model is the multicolored Asian lady beetle, *Harmonia axyridis*, which displays significant elytral spot and color polymorphism. Using transcriptome data from two life stages, adult and larva, we characterize the transcriptome, thereby laying a foundation for further analysis and identification of the genes responsible for the continual maintenance of spot variation in *H. axyridis*.

## Introduction

The evolution and maintenance of phenotypic polymorphism and striking visual phenotypes have fascinated scientists for many years (Darwin, 1859; Endler, 1986; Fisher, 1930; Gray and McKinnon, 2006; Joron et al., 2006). In general, insects have become increasingly popular as study organisms to examine phenotypic variation (Jennings, 2011; Joron et al., 2006). One such insect displaying extensive elytra and spot variation that has yet to be extensively studied is the Asian Multicolored Ladybeetle, *Harmonia axyridis*.

The mechanisms responsible for the evolution of these phenotypes are as widespread as the species that display them. Aposematism, crypsis, and mimicry may play a role in the evolution of phenotypic variation in the animal kingdom. Members of family *Dendrobatidae*, poison dart frogs, are aposematically colored (Cadwell, 1996), while *Tetrix subulata* grasshoppers maintain their phenotypic polymorphism to aid in crypsis (Karpestam et al., 2014). A mimicry strategy is utilized by one particularly well-characterized species that exhibits phenotypic polymorphism, the Neotropic butterfly system, *Heliconius*. The color, pattern, and eyespot polymorphism seen in *Heliconius* is thought to have arose as a result of Müllerian mimicry (Flanagan et al., 2004) and the supergenes underlying these traits have been well characterized (Kronforst et al., 2006, Joron et al., 2006, Jones et al., 2012).

These studies, aiming to elucidate the mechanistic links between phenotype and genotype, present a unique opportunity to gain insight into the inner workings of many important evolutionary processes. While systems like poison frogs and butterflies have been pioneering, the use of novel models, especially those that can be easily manipulated, are needed. One such study system that possesses many of the benefits of classical models, while offering several key benefits is the multicolored Asian lady beetle, *Harmonia axyridis. Harmonia*, which is common throughout North America, and easily bred in laboratory environments, possesses significant variation in elytral spot number and color.

Elytra color can be red, orange, yellow, or black and spot numbers of *H. axyridis* range from zero to twenty-two (personal observation). The patterning is symmetrical on both wings. In some animals, there is a center spot beneath the pronotum which leads to an odd number of spots. The elytral spots are formed by the production of melanin pigments (Bezzerides et al., 2007). The frequency of different morphs varies with location and temperature. The melanic morph is more prevalent in Asia when compared to North America (LaMana and Miller, 1996; Dobzhansky, 1993). A decrease in melanic *H. axyridis* has been shown to be correlated with an increase in average yearly temperatures in the Netherlands (Brakefield and de Jong, 2011).

Sexual selection may play a role in color variation in *H. axyridis*. Osawa and Nishida (1992) remarked that female *H. axyridis* might choose their mates based on melanin concentration. Their choice, however, has been shown to vary based on season and temperature. Non-melanic (red, orange, or yellow with any spot number) males have a higher frequency of mating in the spring-time, while melanic (black) males have an increased frequency of mating in the summer. While this has been shown with respect to elytral color, no such findings have occurred for spot number. Although these spot patterns are believed to be related to predator avoidance, thermotolerance, or mate choice (Osawa and Nishida, 1992), the genetics underlying these patterns is currently unknown.

To begin to understand the genomics of elytral coloration and spot patterning, we sequenced the transcriptome of an late-stage larva and adult ladybug. These results lay the groundwork for future study of the genomic architecture of pigment placement and development in *H. axyridis*.

### Methods and Materials

#### Specimen capture, RNA extraction, library prep and sequencing

One larval (Figure 1a) and one adult (Figure 1b) *H. axyridis* were captured on the University of New Hampshire campus in Durham, New Hampshire (43.1339° N, 70.9264° W). The adult was orange with 18 spots. The insects were immediately placed in RNAlater and stored in a −80C freezer until RNA extraction was performed. The RNA from both individuals was extracted following the TRIzol extraction protocol (Invitrogen, Carlsbad USA). The entire insect was used for the RNA extraction protocol. The quantity and quality of extracted RNA was analyzed using a Qubit (Life Technology, Carlsbad USA) as well as a Tapestation 2200 (Agilent technologies, Palo Alto USA) prior to library construction. Following verification, RNA libraries were constructed for both samples following the TruSeq stranded RNA prep kit (Illumina, San Diego USA). To allow multiple samples to be run in one lane, a unique adapter was added to each sample. These samples were then pooled in equimolar quantities. The libraries were then sent to the New York Genome Center (New York, USA) for sequencing on a single lane (125bp paired end) of the HiSeq 2500 sequencer.

**Figure 1a:**
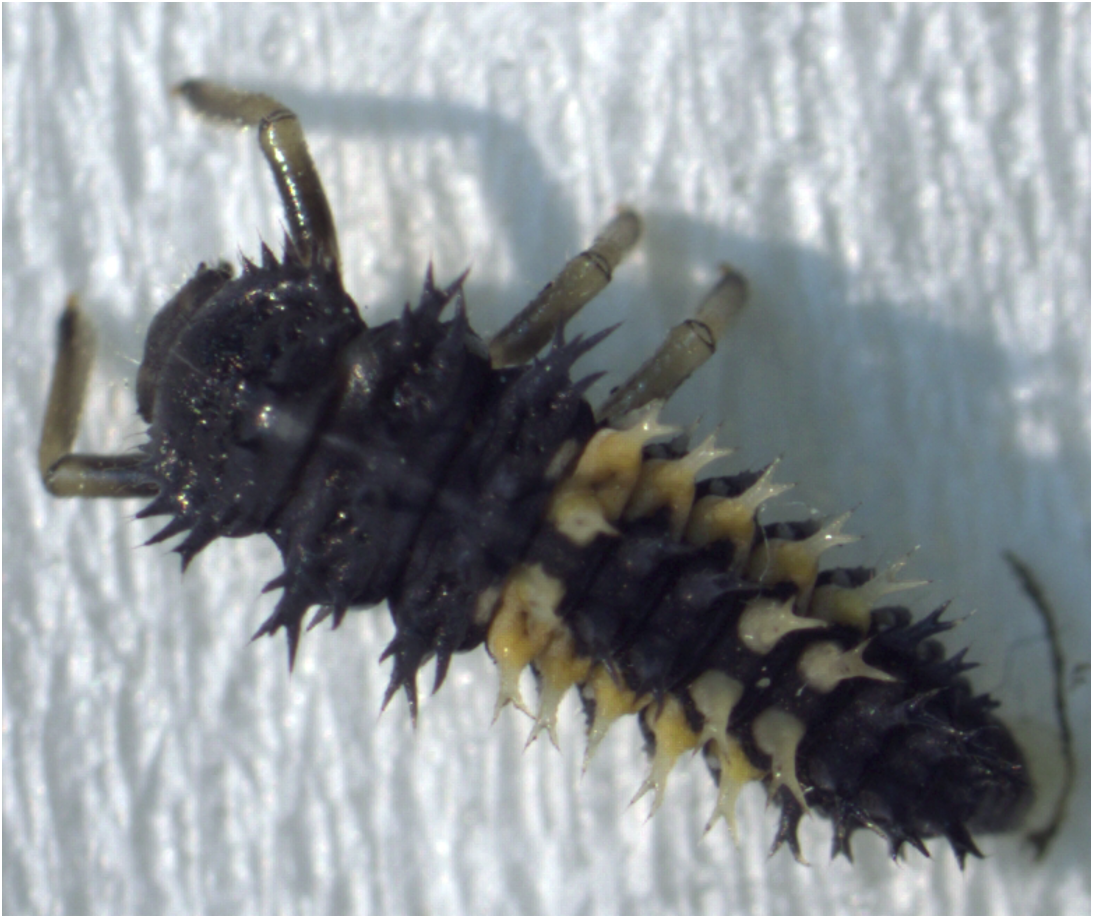
The larva used for transcriptome sequencing.

**Figure 1b:**
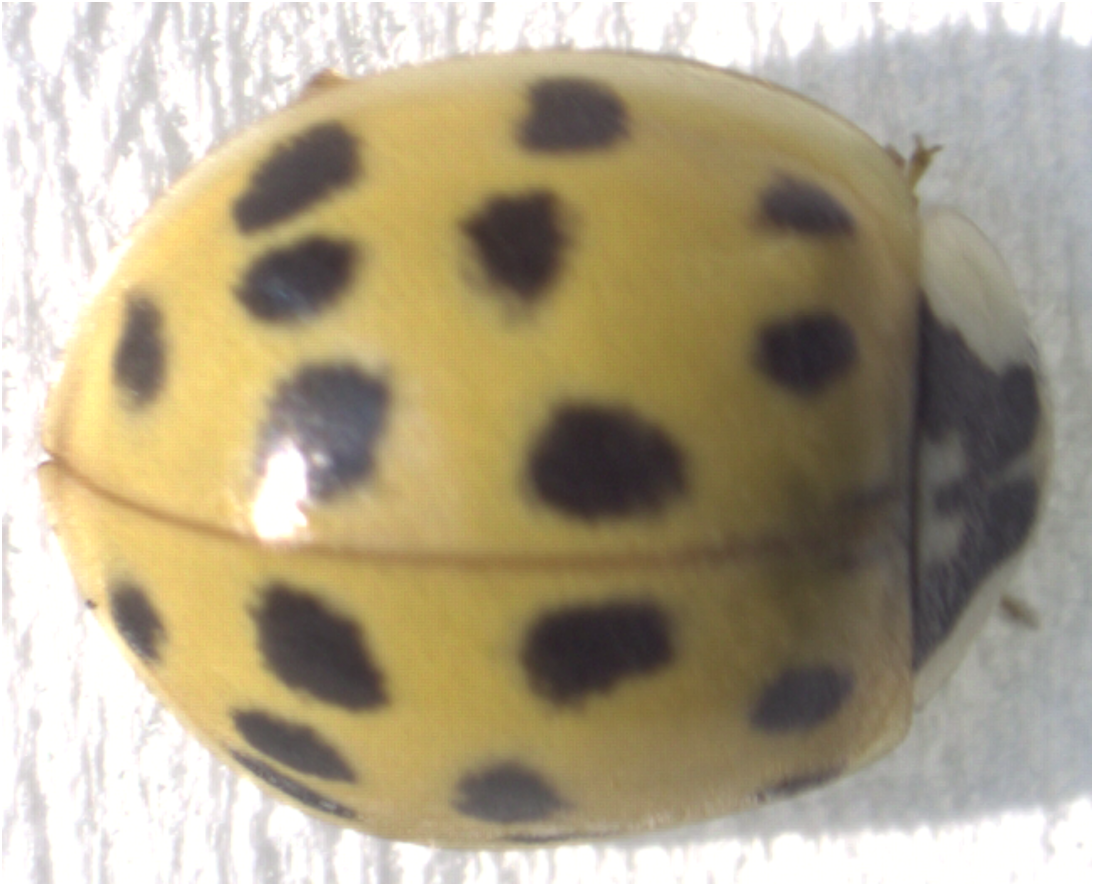
The adult used for transcriptome sequencing.

#### Sequence Data Preprocessing and Assembly

The raw sequence reads corresponding to the two tissue types were error corrected using the software BLESS (Heo et al., 2014) version 0.17 (https://goo.gl/YHxlzI, https://goo.gl/vBh7Pg). The error-corrected sequence reads were adapter and quality trimmed following recommendations from MacManes (MacManes, 2014) and Mbandi (Mbandi, 2014). Specifically, adapter sequence contamination and low quality nucleotides (defined as Phred<2) were removed using the program

Trimmomatic version 0.32 (Bolger, 2014) called from within the Trinity assembler version 2.1.1 (Haas, 2013). Reads from each tissue were assembled together to created a joint assembly of adult and larva transcripts using a Linux workstation with 64 cores and 1Tb RAM. We used flags to indicate the stranded nature of sequencing reads and set the maximum allowable physical distance between read pairs to 999nt (https://goo.gl/ZYP08M).

The quality of the assembly was evaluated using transrate version 1.01 (Unna-Smith 2015; (https://goo.gl/RpdQSU). Transrate generates quality statistics based on a process involving mapping sequence reads back to the assembled transcripts. Transcripts supported by properly mapped reads of a sufficient depth (amongst other things) are judged to be of high quality. In addition to generating quality metrics, transrate produces an alternative assembly with poorly-supported transcripts removed. This improved assembly was used for all downstream analyses and QC procedures. We then evaluated transcriptome completeness via use of the software package BUSCO version 1.1b (Simão 2015). BUSCO searches against a database of highly-conserved single-copy genes in Arthropoda (https://goo.gl/bhTNdr). High quality, complete transcriptomes contain are hypothesized to contain the vast majority of these conserved genes.

To remove assembly artifacts remaining after transrate optimization, we estimated transcript abundance using 2 software packages - Salmon version 0.51 (Patro 2015; https://goo.gl/01UIF6) and Kallisto version 0.42.4 (Bray, 2015; https://goo.gl/BsQMpr). Transcripts whose abundance exceeded 0.5 TPM in either adult or larval datasets using either estimation method were retained. We evaluated transcriptome completeness and quality, again, after TPM filtration, using BUSCO and transrate, to ensure that our filtration processes did not significantly effect the biological content of the assembly.

#### Assembled Sequence Annotation

The filtered assemblies were annotated using the software package dammit (https://github.com/camillescott/dammit; https://goo.gl/05MY5i). Dammit coordinates the annotation process, which involves use of blast (Camacho, 2009), TransDecoder (version 2.0.1, http://transdecoder.github.io/), and hmmer version 3.1b1 (Wheeler 2013). In addition to this, putative secretory proteins were identified using the software signalP, version 4.1c (https://goo.gl/FaOQSj).

To identify patterns of gene expression unique to each life stage, we used the expression data as per above. We identified transcripts expressed in one stage but not the other, and cases where expression occurred in both life stages. The Uniprot ID was identified for each of these transcripts using a blastx search (https://goo.gl/J9saMj), and these terms were used in the web interface Amigo (Carbon et al., 2009) to identify Gene Ontology terms that were enriched in either adult or larva relative to the background patterns of expression. The number of unique genes contained in the joint assembly was calculated via a BLAST search of the complete gene sets of *Homo sapiens, Drosophila melanogaster*, and *Tribolium casteneda*.

### Results and Discussion

#### Data Availability

All read data are available via https://goo.gl/VNuHXC. Assemblies and data matrices are available at https://goo.gl/D3xh65.

#### RNA extraction, Assembly and Evaluation

RNA was extracted from whole bodies of both the adult and the larva stage of a single *Harmonia axyridis*. The quality was verified using a Tapestation 2200 as well as a Qubit. The initial concentration for the larva sample was 83.2 ng/uL, while the initial concentration for the adult sample was 74.7 ng/uL. The number of strand-specific paired end reads contained in the adult and larva libraries were 58 million and 67 million, respectively. The reads were 125 base pairs in length.

The raw Trinity assembly of the larval and adult reads resulted in a total of 171,117 contigs (82Mb) exceeding 200nt in length. This assembly was evaluated using Transrate, producing an initial score of 0.10543, and and optimized score of 0.29729. This transrate optimized assembly (89,305 transcripts, 62Mb) was further filtered by removing transcripts whose expression was less than 0.5 TPM. After filtration, 33,648 transcripts (40Mb) remained. To assess for the inadvertent loss of valid transcripts, we ran BUSCO before and after this filtration procedure. The percent of Arthropoda BUSCO’s missing from the assembly rose slightly, from 18% to 21%. Transrate was run once again, and resulted in a final assembly score of 0.29112. This score is indicative of a high-quality transcriptome appropriate for further study. In an attempt at understanding how many distinct genes our transcriptome contained, we conducted a blast search against *Homo sapiens, Drosophila melanogaster*, and *Tribolium casteneda*. This search resulted in 7,246, 7,739, and 7,741 unique matches, which serve as estimates of the number of unique genes expressed in these two life stages. The final assembly is available at https://goo.gl/nWdBuv

#### Annotation

The assembled transcripts were annotated using the software package dammit!, which provided annotations for 23,304, or 69% of the transcripts (available here: https://goo.gl/gpGXLG). These annotations included putative protein and nucleotide matches, 5- and 3-prime UTRs, as well as start and stop codons. In addition to this, analysis with Transdecoder yielded 14,518 putative protein sequences (available here: https://goo.gl/qVLWwD), which were annotated by 4,139 distinct Pfam protein families, while 176 transcripts were determined to be non coding (ncRNA) based on significant matches to the Rfam database (available here: https://goo.gl/x1n7jC). Lastly, 2,925 proteins (7.8% of total) were determined to to be secretory in nature by the software package signalP (available here: https://goo.gl/z0ra1g).

Annotation of the sequence dataset resulted in the identification of host of transcripts that may be of interest to other researchers including: 43 heat-shock and 8 cold-shock transcripts, 87 homoebox-domain containing transcripts, 122 7-transmembrane-containing (18 GPCR’s) transcripts, 13 solute carriers, 143 ABC-transport-containing transcripts, and 21 OD-S (pheromone-binding) transcripts.

A complement of immune-related genes were discovered as well. These include a single member of the Attacins and Coleoptericins, two TLR-like genes, seven Group 1, and 34 Group 2 C-type lectin Receptors (CLRs). Two CARD-containing Cytoplasmic pattern recognition receptor (CRR) genes were discovered, as were 3 MAP kinase containing transcripts. Finally, 119 RIG-I-like receptors (RLR) were found.

Gene expression was estimated for each transcript for both adult and larva (Figure 2, available here: https://goo.gl/wM3TV7). For all transcripts expressed in the adult, the mean TMP=29.7 (max= 80,016.9, SD= 500.8) while the mean larval TPM=29.71 (max= 166,264, SD=941). When analyzing transcripts found uniquely in these two tissues, mean adult TMP=9.9 (max=3,037, SD=86) and for larva TPM=8.6 (max=687, SD=41).

**Figure 2:**
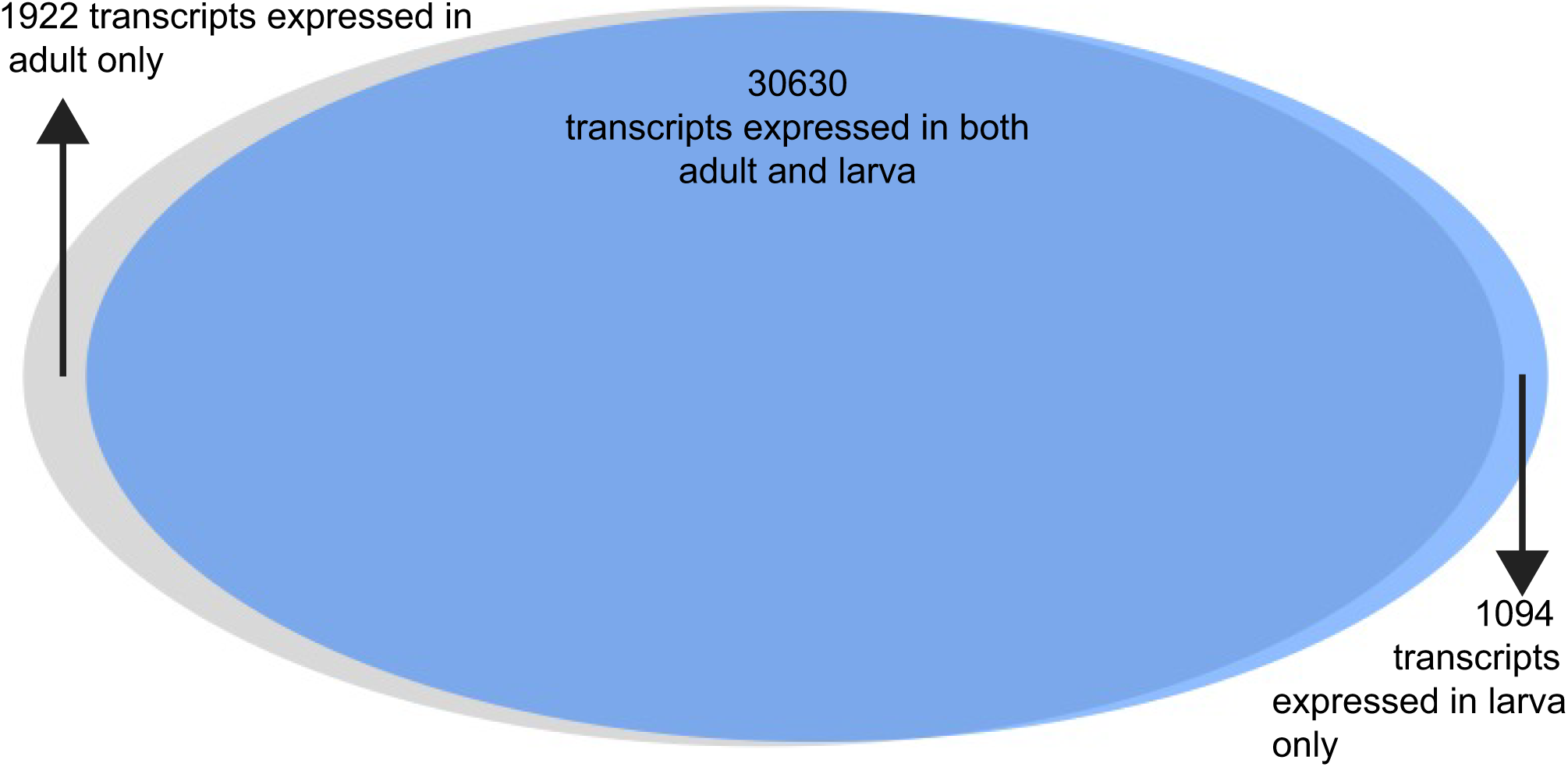
The Venn diagram representing the number of transcripts expressed in both adult and larva, as well as those expressed uniquely in one or the other.

Analysis of the differences between adult and larval life stages were carried out as well. Because these life stages were only sequenced with a single individual each, they should be interpreted with some caution. The vast majority of transcripts were observed in both life stages (n=30,630, 91%), with a small number being expressed uniquely in larva (n=1,094) and adult (n=1,922). Of these transcripts expressed uniquely in either larva or adult, 6.1% and 4.6%, respectively, were found to be secretory in nature.

## Conclusions

Phenotypic polymorphisms and striking visual phenotypes have fascinated scientists for many years. The breadth of evolutionary causes for the maintenance of these phenotypes are as numerous as the species that display them. One organism, *Harmonia axyridis*, provides a unique opportunity to explore the genetic basis behind the maintenance of an easy to quantify variation - elytral spot number. While understanding these genomic mechanisms is beyond the scope of this paper, we do provide a reference transcriptome for *H. axyridis*, a foundational resource for this work.

This study indicates that most gene expression profiles are shared across life stages of *H. axyridis*. While the majority of proteins identified in the assembled transcriptome were structural in function, analyses of protein families using the Pfam database indicated the presence of pigment proteins. In particular, RPE65, which functions in the cleavage of carotenoids, was found. In *H. axyridis*, increased carotenoid pigmentation has been linked to increased alkaloid amounts (Britton et al., 2008). In addition, the elytral coloration of the seven spot ladybug, *Coccinella septempunctata*, is a result of several carotenoids (Britton et al., 2008). While larva are mostly black (**Figure 1a**), we posit that the orange sections on the lower back could be due to carotenoid production. Moreover, this study provides a necessary foundation for the continued study of the genetic link between genes and the maintenance of variation in *H. axyridis*.

